# Hippocampal representations differentiate reactive and anticipatory responses during foraging under threat

**DOI:** 10.64898/2026.04.17.719234

**Authors:** Chelsey C. Damphousse, Olivia L. Calvin, A. David Redish

## Abstract

Adaptive behavior under threat requires balancing reward pursuit against the risk of harm. During approach-avoidance conflict, animals often pause at decision points, but whether these pauses reflect a unified process or distinct decision states remains unclear. Here, we replicate and extend findings from Calvin et al.^1^ by analyzing hippocampal activity in rats performing a pseudo-predator foraging task across two cohorts. We compared mid-track aborts (MTAs), mid-track continues (MTCs), and retreats. Behaviorally, MTAs and MTCs emerged from a shared pause state but diverged in outcome, whereas retreats reflected rapid escape following attack. Despite similar endpoints, retreats and MTAs differed in movement dynamics and neural activity. During retreats, hippocampal representations remained biased toward the attack location. In contrast, representations during MTAs shifted toward safe locations. During pauses, representations differed according to subsequent behavior: pauses preceding MTAs showed greater decoded probability near the attack region, whereas pauses preceding MTCs showed greater decoded probability to goal locations. These differences were present during the outbound approach, suggesting decision-related processes emerged early on in the journey. Together, these findings dissociate hippocampal representations associated with reactive escape from those underlying anticipatory, anxiety-like decision-making, suggesting that the hippocampus dynamically tracks behaviorally relevant features to guide decisions under threat.

## Introduction

Adaptive behavior often requires balancing reward pursuit against the risk of harm. In approach-avoidance conflict, animals must decide whether to continue toward a goal or withdraw in response to potential threat. Such conflicts are a central feature of anxiety-like behavior, in which actions can be guided by uncertainty about potential harm rather than immediate danger ^2–4^. A prominent behavioral feature of approach-avoidance conflict is hesitation: animals frequently pause at locations where threat is anticipated, even while motivated to proceed ^5^. These pauses are often interpreted as reflecting internal evaluation processes, but their functional role and neural basis remain poorly understood.

Hesitation has also been studied in the context of approach-approach conflict in which animals pause at decision points while hippocampal ensembles transiently represent possible future trajectories ^6,7^. These dynamics have been associated with deliberative evaluation in reward-guided decisions. Recent work by Calvin et al.^1^ examined hippocampal activity during foraging in the presence of a robotic predator (adapted from Choi and Kim^8^). In this task, rats ran past a robotic predator in order to obtain a small food reward. On select trials, the robot would attack pseudorandomly when an invisible threshold on the maze was crossed, moving forward into the “threat zone” toward the rat. Following experience with an attack, rats exhibited pauses along the track when approaching the threat zone, and hippocampal population activity during these pauses reflected structured spatial representations related to threat. These findings suggest that the hippocampus may contribute to processing threat-related information during moments of behavioral uncertainty in a similar manner to that during deliberative decisions between multiple positively-valenced outcomes to approach.

Calvin et al.^1^ focused on pause-related activity during approach to threat without directly testing whether pauses that ended in abortive return versus continued approach reflected distinct underlying neural processes. In addition, responses to actual threat encounters — such as rapid retreat following attack — were not examined, leaving open the question of how hippocampal representations differ between anticipatory and reactive defensive behaviors.

Here, we replicate and extend the analysis of Calvin et al.^1^, directly comparing pausing behaviors on outbound journeys — mid-track aborts (MTAs), in which animals terminate their approach and return to the nest, and mid-track continues (MTCs), in which animals resume forward movement after pausing. We further compared retreat behavior following attacks, which reflects a reactive escape response rather than an anticipatory decision, to MTAs, which reflect anticipatory worry about future outcomes. We further extend the work by analyzing behavioral changes during extinction (with the robot present or absent) and re-exposure to attack.

We recorded neural ensembles from the dorsal hippocampus in rats performing the same predator-based foraging task as Calvin et al.^1^ and applied a unified analysis pipeline across cohorts, including reanalysis and extension of the original dataset (cohort 1) and collection of a new cohort (cohort 2). We quantified how spatial representations differed across behavioral epochs and outcomes, and tested whether hippocampal activity distinguishes between MTAs, MTCs, and reactive retreat. These analyses dissociated dorsal hippocampal representations during anticipatory anxiety-like decision-making from those associated with fear-like reactive escape.

## Results

We analyzed two independent cohorts trained on a Gauntlet foraging task (Fig. 1A). Cohort 1 consisted of animals previously reported in Calvin et al.^1^ and is extended here to include previously unanalyzed behavioral data from the extinction phase (EXT) followed by an additional attack session called the re-exposure phase (ReEX). Cohort 2 underwent a similar experimental timeline with minor differences in track and robot configuration. During the linear track (LT) phase, rats shuttled between a nest and a far feeder without threat. In cohort 1, a novelty phase (NOV) introduced the robot without attacks, whereas cohort 2 did not include this phase. During the attack phase (ATTK), the robot lunged across the track on a subset of outbound journeys (cohort 1: 20% probability after 15 “safe” laps; cohort 2: pseudo-randomized schedule after 15 “safe” laps). Attacks occurred exclusively during outbound laps (nest → far feeder; Fig. 1B). Inbound laps were defined as the journey in the opposite direction from the far feeder back to the nest. In the EXT phase, the robot remained present but did not attack for cohort 1 and was removed entirely in cohort 2. Both cohorts concluded with a final single-session attack phase (ReEX).

**Figure 1.**
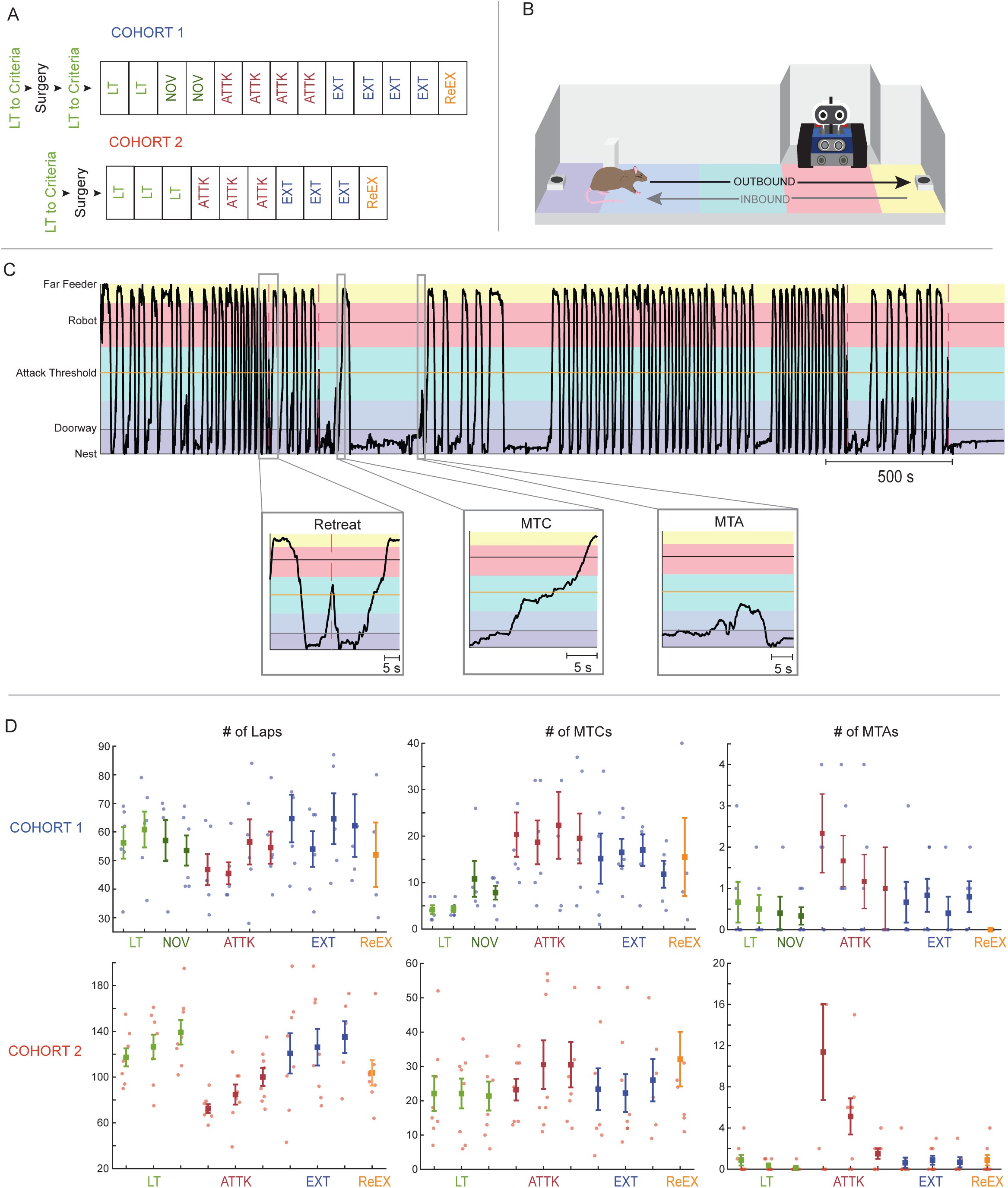
Behaviors replicated between cohorts. a. Experimental timelines for cohort 1 (top) and cohort 2 (bottom). Phases include Linear Track (light green; LT), Novelty (dark green; NOV), Attack (red; ATTK), Extinction (EXT; blue), Re-exposure (orange; ReEX). Each block in the timeline represents one session (1 session per day, 1 hour in length). b. Sideview track schematic when robot is present. Rat runs between nest and far feeder to obtain food rewards, crossing in front of the robot. Attacks only occur during ATTK and ReEX phases, only on outbound journeys (black arrow). Inbound journey shown in grey. For analysis, different regions on the track are shown in different colours, Nest (purple), Track Start (blue), Attack (green), Robot (pink), Far Feeder (yellow) but are not physically present on the actual track. c. Example linearized trajectory on an attack day (example from R823-2023-08-06). Horizontal boxes of different colours represent regions. Landmark positions are shown using solid horizontal lines (doorway in grey, attack threshold in orange, robot in black). Red dashed vertical lines show when attacks occurred. Highlighted sections show behaviors of interest (retreats, MTCs, MTAs). d. Number of laps, MTCs, and MTAs across sessions and phases. Top row: cohort 1 (blue); bottom row: cohort 2 (red). Points represent individual animals, and square markers indicate the mean ± SEM across animals, with marker colour denoting phase. For clarity, extreme outliers are not displayed, but were included in calculation of the error bars. Specifically, 2 outliers were omitted from the laps panel in cohort 1; 2 outliers from the MTC panel in cohort 1 and 3 in cohort 2; and 2 outliers from the MTA panel in cohort 1 and 3 in cohort 2.

### Predator attack evoked three differentiable threat-related behaviors on the track

Predator exposure produced three distinct threat-related behaviors on the track: retreats, MTCs, and MTAs (Fig. 1C). Retreats occurred immediately after a robot attack and consisted of rapid returns toward the nest. In contrast, outbound approaches toward the feeder were frequently interrupted by pauses, particularly after the animals had been attacked once. We focused on these outbound pauses because robot attacks occurred only during outbound travel and pauses were much more common during outbound than return-to-nest journeys. A pause was defined as a cessation of sustained forward movement lasting at least 1 second. The pause epoch then ended when sustained movement resumed, either toward the nest, producing an MTA, or toward the far feeder, producing an MTC. In Calvin et al.^1^, all pauses were described as “on-track pauses.” Here, we distinguish between pauses preceding the decision to abort and return to the nest (mid-track abort, MTA) and those followed by the rat continuing on (mid-track continue, MTC), producing two separable behaviors with a common pause epoch.

In both cohorts, introduction of the attacking robot produced robust changes in behavior (Fig. 1D). Session-level lap counts (Fig. 1D (left)) varied significantly across task phases (linear mixed-effects model; phase effect: F(3,126.06)=26.07, p=3.45×10⁻¹³). In the ATTK sessions, lap completion decreased significantly relative to LT baseline performance in both cohorts (cohort 1: p=0.023; cohort 2: p=2.97×10⁻⁸). Lap counts then increased from ATTK to EXT in both cohorts (cohort 1: p=7.61×10⁻⁴; cohort 2: p=8.55×10⁻⁶). Reintroduction of the attacking robot during ReEX, reduced lap counts relative to LT in cohort 2 (p=0.00135), whereas in cohort 1 this effect was weaker and just reached significance in the planned comparison (p=0.050). Note that differences between cohorts were likely due to continued presence of the robot during EXT in cohort 1 in contrast to its removal in cohort 2.

MTAs showed a strong dependence on task phase (F(3,133)=47.03, p=8.7×10⁻²¹) and increased significantly during the ATTK in both cohorts (cohort 1: p=0.025; cohort 2: p=1.43×10⁻¹⁰; Fig. 1D (right)). MTA counts subsequently declined during EXT (cohort 1: p=0.011; cohort 2: p=1.64×10⁻¹⁰), consistent with the recovery in lap completion. MTCs also showed phase dependence (F(3,133)=12.88, p=1.95×10⁻⁷; Fig. 1D (centre) but differed substantially between cohorts. In cohort 1, MTCs were rare during LT but increased beginning in the NOV phase (p=3.1×10⁻⁴) and remained elevated during ATTK, EXT, and ReEX (all p<10⁻⁵). By contrast, cohort 2 exhibited a relatively high MTC rate during baseline (LT), with further increases during ATTK (p=9.0×10⁻⁵) and ReEX (p=2.0×10⁻⁵). Thus, while MTAs were selectively associated with the attacking robot across both cohorts, MTCs showed cohort-specific differences in baseline prevalence and modulation by phase.

Because both MTAs and MTCs arose from a pause epoch, we next quantified the proportion of pauses that culminated in abortive decisions and asked whether this probability changed within and across phases (Fig. 2A). In cohort 1, abort probability did not vary significantly across ATTK sessions (linear mixed model session effect, *p* = 0.10). However, abort probability was significantly reduced during the ReEX relative to ATTK (*p* = 0.025), indicating that animals did not regain the same likelihood of aborting following exposure to the non-attacking robot over EXT. In contrast, cohort 2 showed a significant change in abort probability across ATTK sessions (session effect, *p* = 0.0069). Although abort probability also declined during ReEX sessions in this cohort, the difference between ATTK and ReEX did not reach statistical significance in cohort 2 (*p* = 0.056). Together, these findings indicate that pauses represented behavioral decision points from which animals either continued toward the feeder or aborted the approach, while also revealing cohort-dependent differences in how abort behavior evolved with experience.

**Figure 2.**
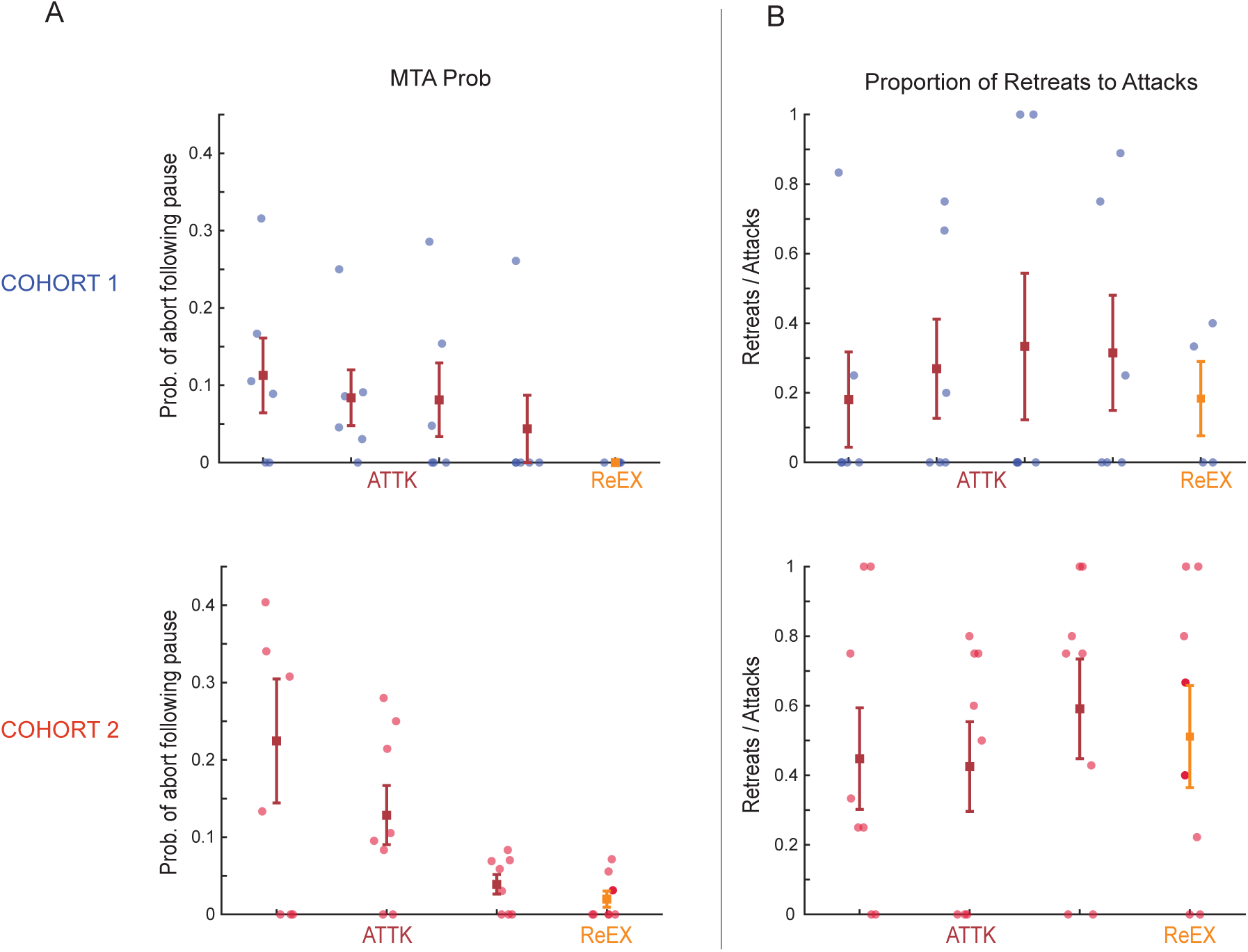
Behavior probability. a. Probability that an outbound pause resulted in an abortive decision (MTA) across sessions in the ATTK and ReEX phases, calculated as MTA / (MTA + MTC). One outlier from cohort 2 is not shown for visual clarity, but was included in the error bar calculation. b. Proportion of attacks that resulted in retreat behavior (retreats / attacks) across sessions of the ATTK and ReEX phase (when robot attacks occurred).

In contrast to MTAs, robot attacks triggered rapid retreats toward the nest without a preceding pause. While both MTAs and retreats resulted in a return-to-the-nest, we differentiated the two based on the external conditions that preceded the behavior — MTAs reflected anticipatory abort decisions made in the absence of an attack, whereas retreats represented a reactive response triggered by an attack. Across sessions, a substantial proportion of attacks resulted in retreat behavior (Fig. 2B). Unlike abort probability, retreat probability remained stable across sessions and phases in both cohorts. In cohort 1, the proportion of attacks that produced retreats did not change significantly across attack sessions (linear mixed model session effect, *p* = 0.083) and did not differ between ATTK and ReEX (*p* = 0.79). Similarly, in cohort 2, retreat probability did not change across ATTK sessions (*p* = 0.23) and did not differ between phases (*p* = 0.83). These findings indicate that retreat behavior likely reflects a consistent reactive escape response rather than a deliberative decision process that evolves with experience.

### Retreats and MTAs differed in movement dynamics

Speed analyses revealed complementary changes in movement dynamics across task phases (Fig. 3). Relative to each rat’s average LT speed, inbound speed (Fig. 3A) increased during ATTK in both cohorts (cohort 1, *p* = 0.00477; cohort 2, *p* = 0.00207), indicating faster returns toward safety under threat. In contrast, outbound speed (Fig. 3B) decreased sharply during ATTK in both cohorts (cohort 1, *p* = 1.29 × 10⁻⁶; cohort 2, *p* = 3.03 × 10⁻¹⁰), consistent with slower, more cautious approaches toward the feeder. During EXT, inbound speed showed partial attenuation: it did not differ from ATTK in cohort 1 (*p* = 0.501), but declined significantly from ATTK in cohort 2 (*p* = 0.00265). Outbound speed, by contrast, partially recovered during EXT in both cohorts (cohort 1, *p* = 0.00269; cohort 2, *p* = 1.92 × 10⁻⁷). During ReEX, inbound speed was elevated relative to LT but did not reach significance in either cohort (cohort 1, *p* = 0.081; cohort 2, *p* = 0.0603), whereas outbound speed remained significantly reduced relative to LT in both cohorts (cohort 1, *p* = 0.00687; cohort 2, *p* = 1.19 × 10⁻⁴). Together, these results indicate that robot attack shifted locomotor behavior in opposite directions, promoting slower outbound approaches toward reward but faster inbound returns to safety.

**Figure 3.**
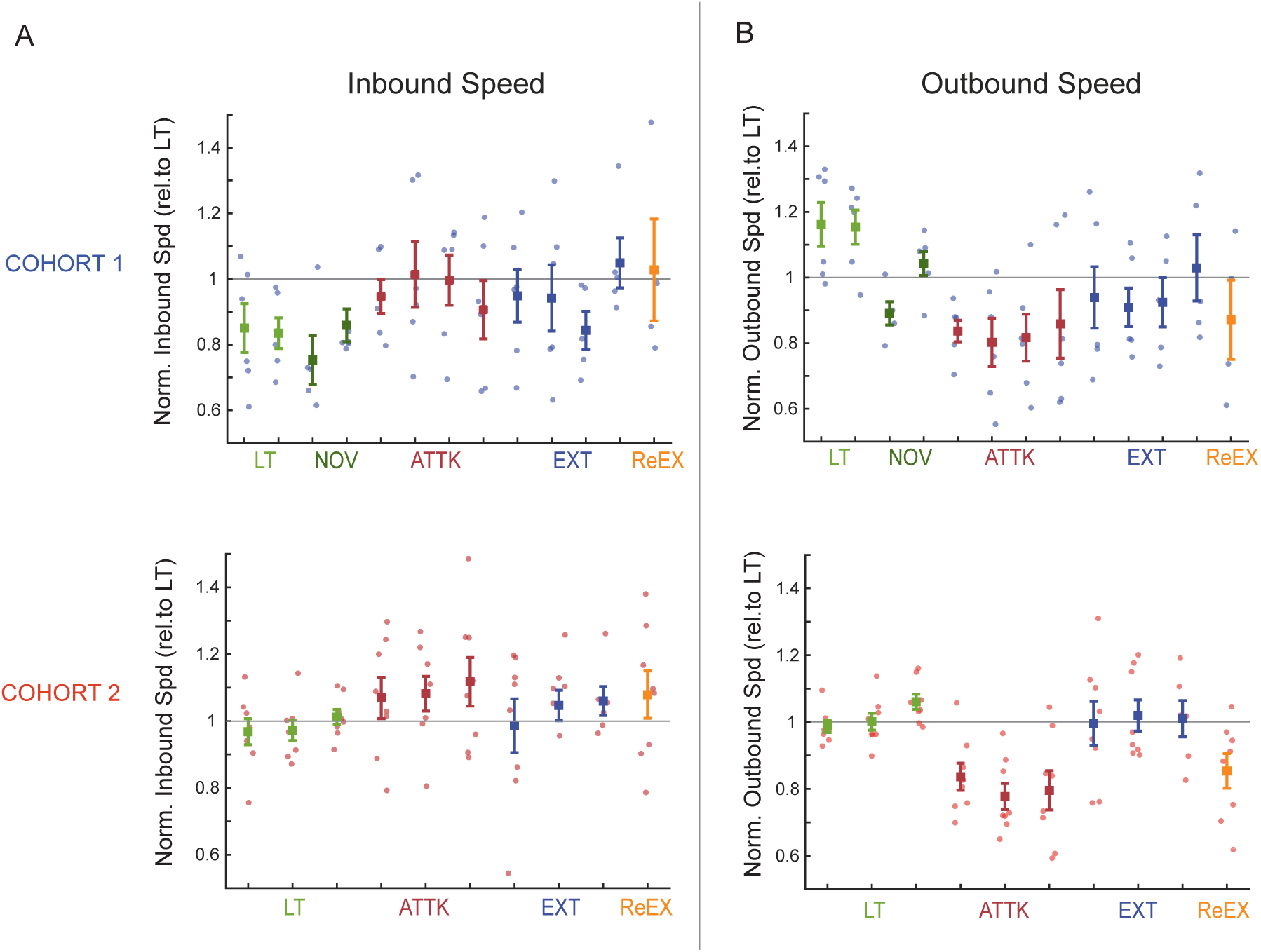
Speed dynamics. a. Relative inbound speed across sessions and phases, relative to linear track (LT) baseline. Top row: cohort 1; bottom row: cohort 2. Points represent individual animals (colour-coded by cohort), with square markers indicating the mean ± SEM across animals (colour-coded by phase). b. Same as left but for outbound speed.

To distinguish retreats from MTAs, we compared inbound speeds for each behavior to matched inbound control laps (Fig. 4). Since all three behaviors only occurred during ATTK, analysis was restricted to this phase. In cohort 1, retreat inbound speeds were significantly higher than control inbound laps (*p* = 2.25 × 10⁻⁴), whereas MTA inbound speeds were significantly lower than controls (*p* = 3.05 × 10⁻¹⁵); retreat inbound speeds were also much faster than MTA inbound speeds (*p* = 3.11 × 10⁻¹²). Cohort 2 showed the same qualitative pattern: retreat inbound speeds exceeded control inbound laps (*p* = 2.22 × 10⁻⁴), MTA inbound speeds were markedly reduced relative to controls (*p* = 3.44 × 10⁻⁴⁸), and retreat inbound speeds were significantly faster than MTA inbound speeds (*p* = 3.40 × 10⁻¹⁹). Across cohorts, threat-related behavior had a strong effect on inbound speed (mixed-effects ANOVA: *F* = 426.26, *p* = 5.25 × 10⁻¹⁰⁹). Thus, while retreats were characterized by rapid flight toward the nest, MTAs were associated with markedly slower inbound movement. This reduction in inbound speed following MTAs dissociates anticipatory abort decisions from reactive retreat behavior, suggesting that abortive returns reflect cautious withdrawal rather than the rapid escape that follows an attack.

**Figure 4.**
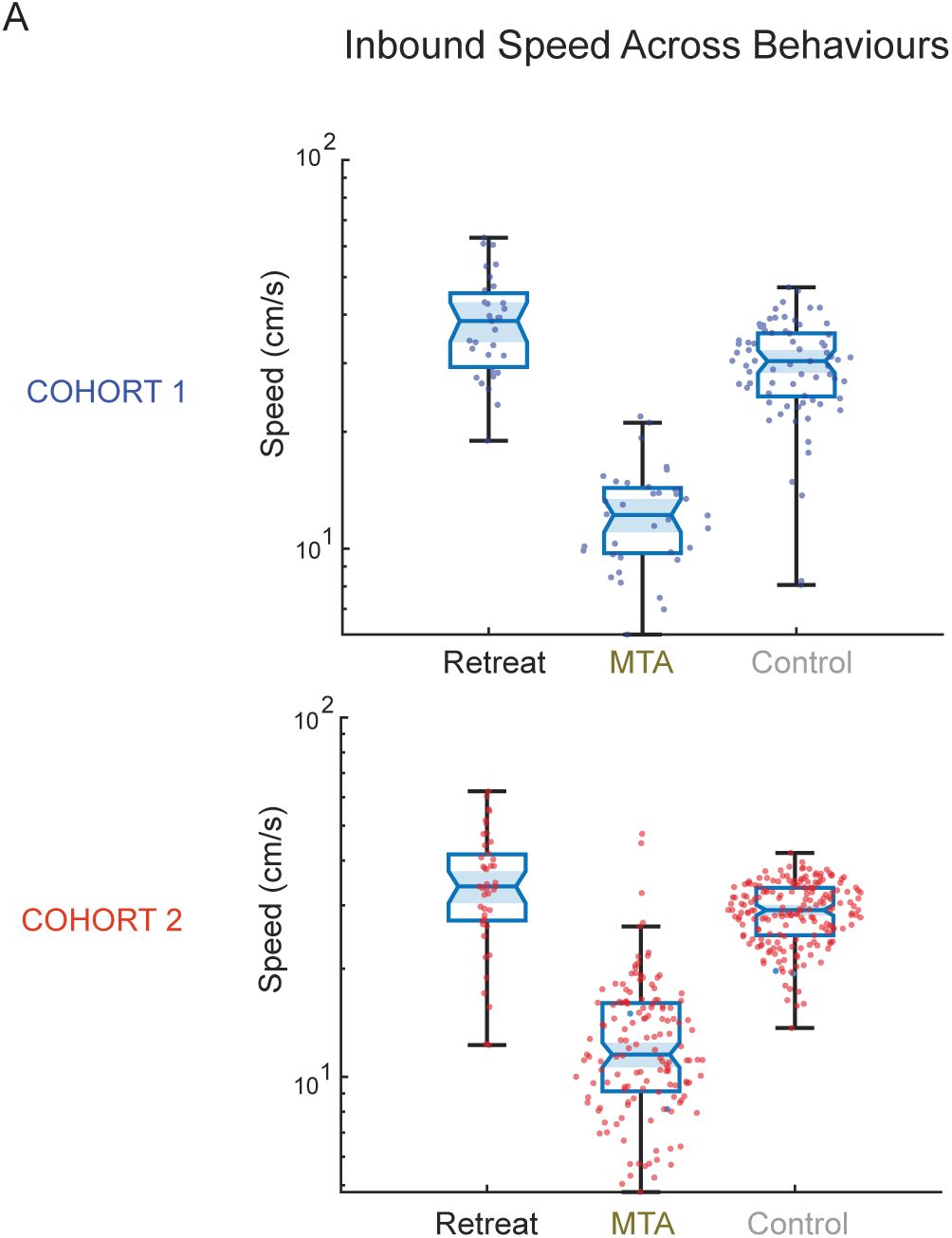
Speed differences between behaviors. a. Mean inbound speed associated with retreats, MTAs, and matched inbound control laps during ATTK. Top row: cohort 1; bottom row: cohort 2.

These behavioral differences raised the question of whether retreats and MTAs were also associated with distinct hippocampal representations.

### Hippocampal representations differentiated retreats from MTAs

Both retreats and MTAs ultimately resulted in the animal turning and fleeing toward the nest. However, these behaviors arose under different conditions: retreats occurred immediately after robot attack, whereas MTAs followed pauses during approach in the absence of an attack (Fig. 5A). To determine whether decoded hippocampal representations differed between these behaviors, we aligned decoded position to the beginning of the inbound journey for retreats and MTAs and compared their decoded spatial representations.

**Figure 5.**
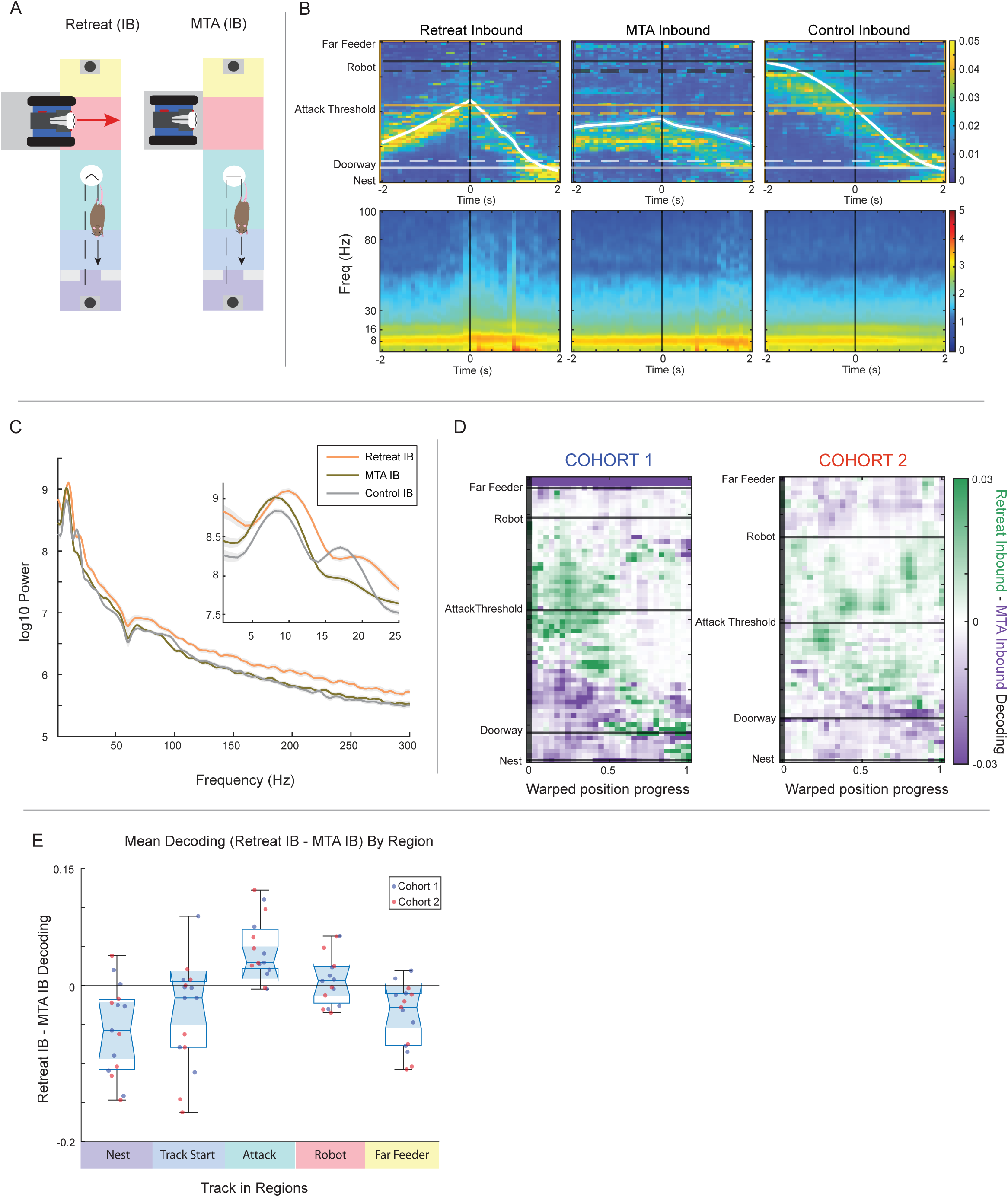
Comparison of inbound journey following retreats and MTAs. a. Aerial-view schematic of inbound journey following a retreat (left) and mid-track abort (MTA; right). Retreats are triggered by robot attack (red arrow), whereas MTAs follow anticipatory pause during approach. Track regions are colour-coded as in Fig. 1. Dashed lines show rat trajectory with curved turnaround representing the rapid change in direction during a retreat and the horizontal line representing a stationary pause during an MTA. Feeders are shown at either end and grey blocks represent the doorway. White circles highlight the turnaround point. b. Top: Decoded position aligned to turnaround during Retreats (left), MTAs (center), and matched inbound control laps (right). Window is shown for 2 seconds on either side of the event. Black vertical line represents the event of interest, in this case the moment inbound travel begins. Horizontal lines show landmark locations (Doorway; light grey, Attack threshold; orange, robot; dark grey). Solid lines represent the landmark position for cohort 1, and dashed for cohort 2. Bottom: Corresponding time-frequency power spectrograms. c. Power spectral density (PSD) curves (1-300 Hz) for Retreat Inbound (IB; orange), MTAs IB (brown), and Control IB (grey), averaged across rats (cohorts combined). Inset shows 1-25Hz range of the same data. d. Difference in decoded position during inbound return (Retreat IB - MTA IB). Positions were spatially warped to account for differing journey lengths. Cohort 1 is shown on the left and cohort 2 on the right. Green indicates stronger representation in the first condition of the subtraction (Retreat IB), whereas purple indicates stronger representation in the second condition (MTA IB). Horizontal lines denote key landmarks. e. Mean decoding bias (Retreat IB - MTA IB) across track regions (cohorts combined). Positive values indicate stronger representation during Retreat IB, whereas negative values indicate stronger representation during MTA IB.

Event-aligned decoding revealed distinct temporal dynamics in hippocampal spatial representations during retreat and MTA inbound journeys. In Fig. 5B, the white trace shows the mean aligned position of the animals, and the black vertical line marks time 0, defined as the moment when the pause stopped and inbound motion began. Warmer colours on the plot indicate higher decoded probability at a given track location in the group-averaged posterior. Shown on the y axis using horizontal lines, actual locations of the attack threshold (not a visually distinct location but rather a position on the track that must be crossed for the robot to potentially attack), robot, feeders and doorway are shown. In the retreat condition (Fig. 5B; left), subjects rapidly turned and moved toward the nest, but the decoded probability remained concentrated above their current position, particularly around the attack-threshold region. Thus, even as they fled inbound, hippocampal representations continued to emphasize the recent danger behind them. In contrast, during MTA inbound journeys (Fig. 5B; centre) decoded probability was initially distributed near the turnaround location but then shifted to lower track positions along the animal’s actual return path toward the doorway and nest. Relative to retreats, MTA inbound decoding was therefore less persistently anchored to the attack region and instead encoded the trajectory back to safety. In contrast to both retreats and MTAs, control inbound journeys (Fig. 5B; right) showed a comparatively smooth correspondence between decoded position and the animal’s ongoing movement.

The event-aligned spectrograms shown in the bottom row of Fig. 5B suggest accompanying differences in oscillatory LFP state. In particular, retreat inbound events showed visibly greater low-frequency power around the alignment point than either MTA inbound events or matched inbound controls, prompting us to summarize these spectral differences across sessions in Fig. 5C. In the session-level PSD curves (Fig. 5C), retreat inbound showed the greatest increase in overall power, with a moderate increase in MTA inbound compared to control journeys. This separation was especially clear in the low-frequency range highlighted in the inset, where retreats exhibited elevated power around ∼4-10 Hz relative to both MTAs and controls. MTAs also showed somewhat greater low-frequency power than controls, but less than retreats. In addition, both retreats and MTAs showed a relative reduction in power around ∼16-18 Hz compared with control inbound journeys.

Because individual retreats and MTAs occurred at diverse locations on the track and thus differed in journey length, we spatially warped the inbound trajectories to enable direct comparison across events (Fig. 5D). In these subtraction plots, green shades indicate positions that were more strongly represented during retreats, whereas purple shades indicate positions that were more strongly represented during MTAs. This normalization revealed a spatial bias in decoding between the two behaviors across both cohorts. In particular, retreats showed stronger decoding around the attack threshold across much of the inbound trajectory. By contrast, MTAs showed stronger representation of locations nearer the nest, as well as more distal nonlocal locations later in the trajectory. To quantify these effects, we divided the track into contiguous behaviorally relevant regions spanning the full track (as shown in Fig. 1B). Quantification of the mean decoding difference (Retreat IB - MTA IB) across track regions revealed a significant main effect of zone (Fig. 5E; ANOVA: p = 2.37 × 10⁻⁶). Post-hoc comparisons identified significant differences in the Nest, Attack, and Far Feeder regions (Bonferroni-corrected α = 0.01), with MTAs biased toward the Nest and Far Feeder locations and retreats biased toward the Attack region.

Together, these analyses support the idea that retreat and MTA inbound journeys were not distinguished simply by different movement kinematics or trajectory lengths, but by distinct patterns of hippocampal representation. Although both behaviors culminated in return-to-nest movement, retreats showed greater decoded probability near the recent site of danger, whereas MTAs showed greater decoded probability toward safe and distal locations. These findings suggest that reactive retreat may reflect continued processing of immediate threat, whereas MTA inbound behavior may reflect a qualitatively different, more anticipatory return state.

### Hippocampal representations during pauses differentiated abort from continue decisions

In addition to retreats triggered by attacks, animals frequently paused during outbound approaches before either aborting the approach (MTA) or continuing toward the feeder (MTC; Fig. 6A). Because the pausing epochs were similar between the two behaviors and preceded the behavioral output of aborting or continuing, the pause period provided an opportunity to test whether hippocampal activity during the pause reflected the upcoming behavioral decision. To provide an easy means of comparison, the analysis for MTA Pause vs. MTC Pause (Fig. 6) follows the same layout as the analysis for Retreat Inbound vs. MTA Inbound (Fig. 5).

**Figure 6.**
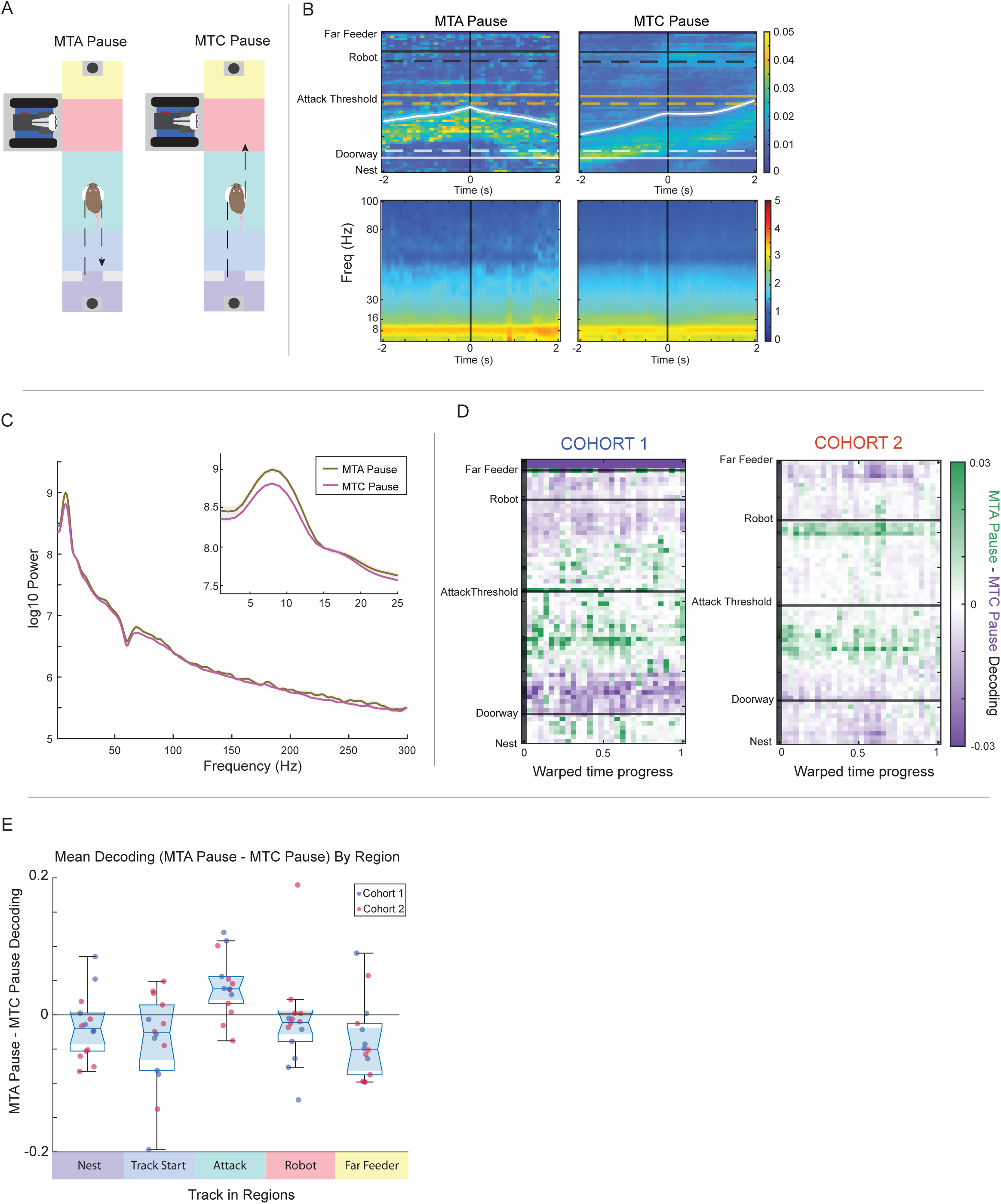
Comparison of pauses preceding abort and continue. a. Schematic of pause preceding an abort (MTA; left) and continuing on to the far feeder (MTC; right). Track regions are colour-coded as in Fig. 1. Dashed lines show rat trajectory. Feeders are shown at either end and grey blocks represent the doorway. White circles highlight pause period. b. Top: Decoded position aligned to middle of pause period in MTAs (left), MTCs (right). Window is shown for 2 seconds on either side of the event. Black vertical line represents the event of interest. Horizontal lines show landmark locations for cohort 1 (solid), and cohort 2 (dashed). Bottom: Corresponding time-frequency power spectrograms. c. Power spectral density (PSD) curves (1-300 Hz) for MTA pause (brown), MTC pause (pink), averaged across rats (cohorts combined). Inset shows 1-25Hz range of the same data. d. Difference in decoded position during pause period (MTA Pause - MTC Pause). Positions were temporally warped to account for differing pause durations. Cohort 1 is shown on the left and cohort 2 on the right. Green indicates stronger representation during the MTA pause, whereas purple indicates stronger representation during the MTC pause. Horizontal lines denote key landmarks. e. Mean decoding bias (MTA Pause - MTC Pause) across track regions (cohorts combined). Positive values indicate stronger representation during the MTA Pause, whereas negative values indicate stronger representation during the MTC Pause.

Decoded position aligned to pause onset revealed differences in hippocampal spatial representations between pauses that were followed by MTAs versus MTCs (Fig. 6B). Because these plots are aligned to pause onset, rather than pause offset, the +1 second time point often still fell within the pause itself. MTA and MTC pauses were defined as lasting at least 1 second, and many pauses exceeded this minimum duration. Thus, the limited forward displacement visible during MTC pauses shortly after pause onset reflects the ongoing pause period. Decoded position aligned to pause onset revealed distinct representational trajectories preceding the two behavioral outcomes (Fig. 6B). During pauses that preceded MTAs (Fig. 6B; left), decoding remained biased toward locations near the attack region, whereas pauses preceding MTCs (Fig. 6B; right) showed relatively stronger representation of locations ahead of the animal along the prospective outbound trajectory.

The event-aligned spectrograms shown in the bottom row of Fig. 6B suggested only modest differences in oscillatory LFP state between the two behaviors. In particular, MTA pauses appeared to show somewhat stronger low-frequency power around ∼8 Hz than MTC pauses. Summarizing these spectral differences in Fig. 6C likewise revealed broadly similar overall frequency profiles during pause periods of MTAs and MTCs, with only a modest low-frequency elevation during MTA pauses. Thus, compared with the clearer differences in decoded spatial content, pause-related spectral differences were relatively subtle, suggesting that the two behaviors were more strongly differentiated by the spatial content of hippocampal representations than by large changes in global oscillatory LFP state.

Because MTA and MTC pauses differed in duration, we temporally warped pause epochs to account for this difference (Fig. 6D). In these subtraction plots, green shades indicate positions that were more strongly represented during MTA pauses, whereas purple shades indicate positions that were more strongly represented during MTC pauses. Across both cohorts, the most prominent green band was present around the attack threshold, indicating stronger attack-related decoding during pauses that ultimately ended in abortive decisions. Cohort 2 shows a large band of green around the robot location which is not present in cohort 1. Purple values were weaker and less spatially concentrated, suggesting that MTC pauses were comparatively less dominated by threat-related representations.

We next quantified mean decoding bias across predefined track regions (Fig. 6E). This analysis revealed a significant main effect of zone (ANOVA: p = 0.0064), and post-hoc comparisons identified a significant difference specifically in the attack region (Bonferroni-corrected α = 0.01), with pauses preceding MTAs exhibiting greater decoding of the attack region. Together, these findings indicate that the pause already contained a threat-related representational bias that predicted whether the animal would ultimately abort or continue the approach.

### Threat-related hippocampal representations emerged before the behavioral pause

Since MTA and MTC pauses differed in their spatial representations before the behavior itself diverged (Fig. 6), we next asked whether these representational differences were already present earlier during the outbound journey, before the pause occurred (Fig. 7A). For this analysis, the outbound epoch was defined as the pre-pause outbound trajectory: it began when the animal initiated outbound movement from the nest/doorway and ended at pause onset.

**Figure 7.**
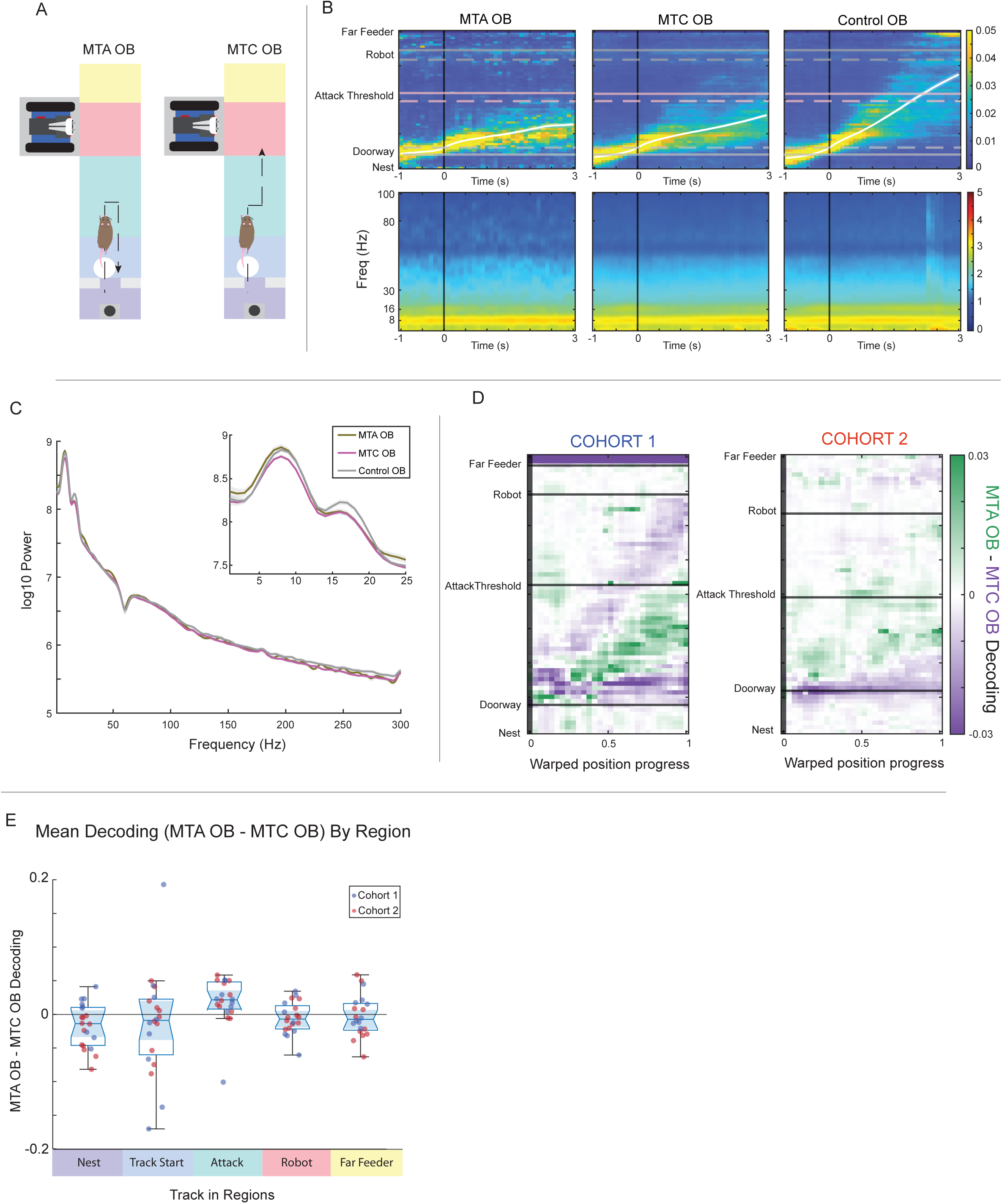
Comparison of outbound journeys preceding pause. a. Schematic of outbound journey preceding the pause associated with an MTA (left) and MTC (right). Track regions are colour-coded as in Fig. 1. Dashed lines show rat trajectory. Feeders are shown at either end and grey blocks represent the doorway. White circles outbound epoch of interest. b. Top: Decoded position aligned to beginning of outbound journey associated with MTAs (left), MTCs (middle), and control laps (right). Window is shown for 1 second before doorway crossing to 3 seconds after. Black vertical line represents the event of interest. Horizontal lines show landmark locations for cohort 1 (solid), and cohort 2 (dashed). Bottom: Corresponding time-frequency power spectrograms. c. Power spectral density (PSD) curves (1-300 Hz) for MTA outbound (OB; brown), MTC OB (pink), and control OB laps (grey) averaged across rats (cohorts combined). Inset shows 1-25Hz range of the same data. d. Difference in decoded position during outbound journey (MTA OB - MTC OB). Positions were spatially warped to account for differing journey lengths. Cohort 1 is shown on the left and cohort 2 on the right. Green indicates stronger representation during the MTA OB, whereas purple indicates stronger representation during the MTC OB. Horizontal lines denote key landmarks. e. Mean decoding bias (MTA OB - MTC OB) across track regions (cohorts combined). Positive values indicate stronger representation during the MTA OB, whereas negative values indicate stronger representation during the MTC OB.

Decoded position aligned to outbound start revealed differences in representational content early in the approach trajectory (Fig. 7B). Here, windows were aligned to the start of outbound movement and shown from 1 s before to 3 s after doorway crossing, rather than using a symmetric ±2 s window, in order to better capture activity emerging once the outbound journey had been initiated. In the MTA outbound condition (Fig. 7B, left), decoding was biased ahead of the animal, particularly toward the attack-threshold region and occasionally extending toward more distal locations. By contrast, MTC outbound decoding (Fig. 7B, centre) more closely followed the animal’s actual outward progression and appeared broadly similar to control outbound laps (Fig. 7B, right). Thus, trajectories that ultimately ended in abortive decisions already showed a stronger nonlocal bias toward threat-related locations at the beginning of the outbound approach.

The event-aligned spectrograms in the bottom row of Fig. 7B suggested only modest oscillatory differences between MTA and MTC outbound trajectories. Both outbound conditions appeared broadly similar overall, although each showed a relative reduction around ∼16 Hz compared with control outbound laps. This impression was reflected in the session-level PSD summary (Fig. 7C), where MTA and MTC outbound spectra largely overlapped one another and were more similar to each other than either was to control outbound laps.

To directly compare outbound representations between behavioral outcomes, we next spatially warped outbound trajectories to account for differences in journey length (Fig. 7D). In these subtraction plots, green shades indicate positions that were more strongly represented during MTA outbound trajectories, whereas purple shades indicate positions that were more strongly represented during MTC outbound trajectories. Across both cohorts, a strong green cluster was centered on the attack threshold early in the outbound journey, indicating a stronger nonlocal bias toward the threat zone during trajectories that would later terminate in abortive pauses. This bias was not comparably evident in MTC outbound trajectories.

We next quantified mean decoding bias across predefined track regions (Fig. 7E). Although the region-based analysis did not reveal a significant main effect of zone (ANOVA: p = 0.109), post-hoc comparisons identified a significant difference in the Attack region (Bonferroni-corrected α = 0.01; p = 0.00319), with outbound trajectories preceding MTAs showing stronger attack-region representation than those preceding MTCs. Together with the pause-epoch analyses in Fig. 6, these findings indicate that the representational bias associated with abortive behavior emerged before the pause itself, suggesting that hippocampal representations of threat were already evolving during the outbound approach and may have contributed to the subsequent decision to abort.

## Discussion

In natural environments, animals are constantly faced with things to approach and things to avoid. Necessities like food, shelter, and mating opportunities must be pursued, but doing so often requires leaving safe locations and venturing into environments of higher uncertainty. While journeying beyond the nest presents many opportunities for desirable outcomes, it also comes with a risk of undesirable outcomes like predation. Therefore, a central problem is not simply how to approach rewards or avoid threats in isolation, but how to resolve the conflict when presented with a context that contains both ^9–11^. Deliberation between options and outcomes has been studied extensively within approach-approach conflicts ^7,12^, but less between approach-avoidance conflicts. A large body of work has shown that the hippocampus can transiently represent possible future options, and that during deliberative behavior it can simulate alternative trajectories or outcomes before action is taken ^6,13–15^. In approach-approach conflict, hippocampal activity has been shown to reflect prospective evaluation between multiple potential reward-directed paths, supporting the idea that the hippocampus contributes to choice by representing imagined possibilities in a spatial framework ^6,13,16–18^. At the same time, hippocampal representations are not limited to positively valenced locations. Work in both avoidance learning ^19,20^ and threat-related tasks ^1,21^ has shown that the hippocampus can also represent places associated with danger, such as shock zones or other locations to be avoided. What remains less clear is how these representational functions are engaged when an animal must evaluate options of different valence: something to approach and something to avoid, known as approach-avoid conflict ^22–24^.

Addressing how the brain resolves approach-avoid conflicts has increasingly required a shift in how such problems are modeled experimentally. There is growing recognition that understanding adaptive behavior depends on considering the animal’s natural ecology, rather than relying exclusively on classical paradigms ^25–28^. Traditional fear-conditioning tasks often constrain the animal to a narrow set of responses and may therefore provide only a limited view of how the brain organizes and executes behavior under threat. An animal is unlikely to encounter an electrified floor in the wild, but by contrast, it must routinely solve how to forage in the presence of risk and uncertainty. Even studies of hippocampal function differentiating between reward and threat have been typically examined in separable conditions ^20,29^. Foraging offers the possibility of reward, but also the possibility of predation ^9,30^. This tension has motivated the development of robotic-predator paradigms, which place animals in a more ethologically relevant form of approach-avoidance conflict ^1,8,9,21,30,31^. These tasks are powerful not only because they capture a challenge animals are likely to face in nature, but also because they reveal a much richer behavioral repertoire than is typically available in more constrained fear assays. Rather than being placed in a context in which the only way to act is to freeze, animals confronted with a robotic threat during foraging can pause, hesitate, flee, continue approaching, or rapidly retreat following attack. Such paradigms therefore provide a stronger opportunity to ask how neural activity differentiates among multiple defensive and decision-related states, rather than simply indexing the presence or absence of fear.

The behaviors observed on robotic-predator tasks can be understood within the Predatory Imminence Continuum, in which defensive states are organized by an animal’s spatial, temporal, and psychological proximity to threat ^10,32–35^. Within this framework, distal or uncertain threat elicits anticipatory and risk-assessment behaviors, whereas proximal threat triggers rapid, reactive defensive responses ^8,9,21^. Behaviors observed in the present Gauntlet task, as well as in earlier work by Walters et al.^21^ and Calvin et al.^1^, fit well within this organization of defensive modes. Retreats triggered by robot attacks reflect reactive, circa-strike responses to imminent danger, as evidenced by their very fast, ballistic responses. This distinction is consistent with work in mice showing that reactive escape is governed by threat-intensity-dependent threshold mechanisms that rapidly convert sensory threat information into escape initiation ^36^, in contrast to the slower, anticipatory evaluation that appears to characterize MTAs in the present task. Thus, the behavioral structure of the Gauntlet task allows reactive escape to be distinguished from anticipatory decision-making under threat. This makes robotic predator tasks (Gauntlet and Robogator in particular) well suited for asking whether hippocampal representations differ not only between reward and threat, but across different defensive modes that arise as threat becomes more immediate.

To begin to assess hippocampal representations during approach-avoidance conflict, it is useful to consider how options are evaluated in more traditional approach-approach settings. In those tasks, deliberation typically unfolds between two spatially distinct reward locations, often positioned at different arms of a maze. This spatial separation makes it possible to observe hippocampal representations shifting between alternative prospective outcomes. Applying this logic to approach-avoidance conflict requires a task in which the relevant options are similarly dissociable in space. In the classic Choi & Kim Robogator task^8^, the rat approaches a food item positioned between the nest and a robot predator, which is ideal for measuring risky foraging but provides limited spatial separation between reward pursuit and predator location. In the Gauntlet version used here, and in Calvin et al.^1^, the rat must instead run past the robot (*running the Gauntlet*) to obtain reward, separating the location of the predator from the reward site. The inclusion of an attack threshold to be crossed further dissociates the physical location of the robot from the location at which attack is triggered. This spatial decomposition makes it possible to ask what the hippocampus represents at different moments of the conflict: locations to approach, locations to avoid, and the transitions between them.

Having established this spatial structure, we next asked whether hippocampal representations differed across the distinct behavioral resolutions of approach-avoidance conflict. To begin to address this, we extended Calvin et al.^1^ in two important ways. First, we included retreat events, allowing direct comparison between reactive escape following attack and anticipatory return-to-nest behavior in the absence of attack. Second, rather than treating pauses during approach as a single class, we distinguished between pauses that culminated in abortive return to the nest (MTAs) and pauses after which the animal resumed forward movement toward reward (MTCs). These comparisons created two particularly informative contrasts. Comparing retreats and MTAs allowed us to examine a common behavioral endpoint — return to the nest — arising from two very different circumstances: immediate attack versus anticipatory, anxiety-like withdrawal. Comparing MTAs and MTCs, in turn, allowed us to examine a common behavioral interruption — pausing during approach — that resolved into two different outcomes: aborting the approach or continuing on toward reward.

We therefore asked whether decoded spatial content differed between retreats and MTAs. Although both behaviors culminated in return-to-nest movement, this similar behavioral endpoint was associated with different patterns of decoded spatial content. During retreats, hippocampal decoded probability remained elevated toward the site of recent danger, even as the animal moved in the opposite direction, toward the nest. MTAs, by contrast, showed greater decoded probability toward safe and distal locations. Within the framework of the Predatory Imminence Continuum, this dissociation is consistent with the idea that retreats may reflect a more reactive, circa-strike defensive state, whereas MTAs may reflect a more anticipatory, anxiety-like withdrawal in which the animal disengages from approach before direct attack occurs. One of the key strengths of the Gauntlet paradigm is that it elicits both proximal and distal defensive behaviors within the same assay, providing a tractable framework for examining how hippocampal representations shift as threat becomes more immediate.

The LFP changes accompanying retreats and MTAs suggest that these behaviors differ not only in the spatial content of hippocampal representations, but also in the broader oscillatory state of the network. Retreats were associated with a pronounced increase in low-frequency power around ∼4-10 Hz, whereas both retreats and MTAs showed reduced power in the ∼16-18 Hz range relative to matched control inbound journeys. One interpretation is that these effects reflect a transition from routine, ongoing navigation into defensive withdrawal, with retreats representing the most acute version of that state change. The low-frequency enhancement during retreats is consistent with prior work linking 4-Hz-range oscillations to defensive states, including freezing and fear-related responding ^37^. The shared reduction around ∼16-18 Hz may indicate a broader shift in processing mode that accompanies withdrawal to safety ^38^. Because this frequency lies within the beta range, the reduction may reflect disengagement from a more deliberative or ongoing navigational state and entry into a more action-oriented defensive mode ^39^. More broadly, the accompanying higher-frequency differences are reminiscent of proposals that distinct oscillatory bands reflect different hippocampal information-processing modes, including shifts between recollective, evaluative, and action-related states ^20^.

The comparison between MTAs and MTCs further clarified what hippocampal representations may contribute during approach-avoidance conflict. Although both behaviors passed through a common pause state, they diverged in the spatial content expressed during that pause. During pauses preceding MTAs, decoded hippocampal representations showed relatively greater probability near threat-related locations, whereas pauses preceding MTCs showed relatively greater probability ahead of the animal, along the prospective path to reward. This pattern suggests that pausing itself may reflect a common evaluative mode, but that the outcome of that evaluation may depend on what is represented within that state. In this sense, hippocampal activity during the pause may contribute not simply by signaling uncertainty, but by differentially weighting the competing possibilities of withdrawal versus continued approach, consistent with prior work linking hippocampal activity at decision points to deliberative processes and prospective representation of behaviorally relevant outcomes ^6,7,13–15^.

Notably, despite this divergence in representational content, MTA and MTC pauses were not strongly dissociable at the level of overall LFP structure. Their power spectra were broadly similar, with only modest differences. This suggests that the critical distinction between aborting and continuing may lie less in a shift in global hippocampal state and more in the specific spatial content expressed within that state. Pausing may therefore reflect a common evaluative mode, while hippocampal representations within that mode bias behavior toward either withdrawal or continued approach, consistent with evidence that hippocampal ensembles can represent future trajectories prior to overt behavioral commitment ^40^.

Importantly, these representational differences did not emerge only at the pause itself. Outbound trajectories that ultimately culminated in MTAs already showed stronger representation of the attack region than those that led to continued approach. This suggests that the decision to abort was not initiated at the moment of pausing, but may instead have reflected an evolving process that began earlier during approach toward threat. Pauses may therefore be better understood as a behavioral manifestation of an ongoing decision process rather than the point at which that process begins. This interpretation is consistent with prior work showing that hippocampal ensembles can represent future trajectories before overt behavioral commitment ^13–15^, and with evidence that hippocampal dynamics can encode distant, behaviorally relevant locations during successful avoidance ^20^. In this context, our findings suggest that, during approach-avoidance conflict, hippocampal representations are not only prospective, but may be systematically biased toward either threat-related or goal-related locations in a manner that predicts behavioral outcome.

One of the most important implications of the present study is that the hippocampus may represent differently valenced future possibilities within a common deliberative framework. In the approach-approach literature, hippocampal representations have often been interpreted as reflecting possible future choices between multiple positively valenced options ^6,13,15^. The present findings suggest that this framework extends to approach-avoidance conflict: the hippocampus appears not only to represent goals to pursue, but also locations to avoid. In that sense, hippocampal representations may provide a common spatial format for comparing alternatives of different valence. Rather than being limited to reward-directed deliberation, hippocampal prospective activity may support conflict resolution more generally by representing both desirable and dangerous outcomes within the same decision space. This helps reconcile work on hippocampal involvement in memory, spatial cognition, and conflict processing, suggesting that these functions are not separate, but may instead reflect different uses of a shared representational system for guiding behavior under competing demands ^15,22^.

Importantly, these results are correlational and the actual causal mechanism underlying these behaviors likely depend on interactions with other neural circuits such as amygdala ^8,41,42^, accumbens shell ^42,43^, and periaqueductal gray ^44,45^. The observed representational biases between Retreat, MTA, and MTC behaviors likely operate within broader circuits that encode threat, value, and action selection. Work on the original Robogator task demonstrated that amygdala manipulations bidirectionally alter risky foraging, and subsequent recording studies emphasized that amygdala activity in this setting is tightly related to behavioral output ^8^. More recent work has also shown that dopamine signaling in the tail of the striatum promotes avoidance under threat-reward conflict, even at the expense of reward acquisition ^46^. Together, these findings suggest that approach and avoidance behaviors emerge from interacting neural systems rather than a single unified pathway. In this context, hippocampal representations may provide a spatially structured substrate through which competing influences — threat, reward, and action tendency — are evaluated and integrated.

Together, these findings suggest that the hippocampus contributes to decision-making under threat not simply by encoding space or retrieving past experience, but by dynamically representing competing behavioral outcomes across levels of predatory imminence. These representations distinguish reactive and anticipatory defensive states, and bias behavior toward either withdrawal or continued approach. By biasing representation toward either threat or reward before action is taken, hippocampal activity may provide a mechanism for linking internal evaluation to adaptive behavior in environments where survival depends on balancing risk and reward.

## Methods

### Subjects

Two cohorts of Brown Norway rats were used in this study. Cohort 1 consisted of six rats (3 male, 3 female; 7-10 months old) previously reported in Calvin et al.^1^. Cohort 2 consisted of eight rats (4 male, 4 female; 4-10 months old). Animals in cohort 1 were maintained on a 14 h:10 h light-dark cycle, whereas cohort 2 animals were maintained on a 12 h:12 h light-dark cycle.

All rats were food restricted such that their daily caloric intake was earned in the foraging arena via 45 mg nutritionally complete pellets (full nutrition, Test Diet), while remaining above 80% of their free-feeding body weight. Water was available ad libitum in the home cage. All procedures were approved by the University of Minnesota Institutional Animal Care and Use Committee and conducted in accordance with NIH guidelines.

### Surgery

Surgical procedures for cohort 2 were identical to those described in Calvin et al.^1^ except as noted. Rats were initially anesthetized in an induction chamber with 0.5-2% isoflurane in medical-grade oxygen and maintained at this level throughout surgery via a Somnosuite system (Kent Scientific, Torrington, Connecticut, USA). Once deeply anesthetized, the rat was secured in a stereotaxic frame (Kopf Instruments, Tujunga, California, USA), and the skull and dura overlying the dorsal hippocampus were removed. Silicon probes were chronically implanted at an anterior-posterior coordinate of -3.8 mm and a medial-lateral coordinate of ± 2.5 mm (probe details below). The probe and amplifier assembly were affixed to the skull with MetaBond (Parkell, Edgewood, New York, USA), the craniotomy sealed with bone wax, and the entire implant enclosed in a 3D-printed protective shroud (Formlabs, Somerville, Massachusetts).

After surgery, rats recovered in an incubator, received 0.8 mL of Children’s Tylenol orally, and were administered enrofloxacin (25 mg/kg) and carprofen (5 mg/kg) subcutaneously on the day of surgery and for 3 days following. The incision site was cleaned daily with betadine for 7 days, and animals had ad libitum access to food and water for the first 3 postoperative days. Beginning on postoperative day 4, rats were once again placed on food restriction and either resumed behavioral training (cohort 1) or began training (cohort 2).

Probe configurations differed between cohorts. Cohort 1 animals were bilaterally implanted with 64-channel, four-shank Cambridge Neurotech P-1 probes targeting the dorsal hippocampus. In cohort 2, P-1 probes were implanted unilaterally, with the hemisphere of implantation counterbalanced across subjects. In cohort 2 animals, stimulating stereotrodes were additionally implanted in the anterior hippocampal commissure (-1.3 A/P, ± 1.2 M/L, insertion depth -4.0 from the surface of cortex).

### Experimental Setup

#### Arena

The foraging arenas differed slightly between cohorts. For cohort 1, the arena measured 111 cm in total length and consisted of a nest, a linear track, and a robot bay arranged in an L-shaped configuration (see ^1^. The nest measured 19 cm in length and 26 cm in width and connected to a 67 cm long, 26 cm wide linear track. A robot bay was positioned at the far end of the track, measuring 22 cm along the track wall and 35 cm in depth.

The foraging arena for cohort 2 measured 124 cm in total length (including nest and track), 27 cm in width, and 46 cm in height, and was constructed from uniform corrugated plastic sheeting. It consisted of a central linear track flanked by two feeders (Med-Associates, Georgia, Vermont), one at the nest end and one at the far end, and a dedicated robot bay positioned 10 cm from the far feeder. The robot bay measured 33 cm long by 23 cm wide. The nest area measured 18 cm long by 27 cm wide and featured partial walls with an open entrance onto the track. When the robot was not in use, a corrugated-plastic guillotine door was slid into place over the robot bay entrance. Each feeder dispensed two 45 mg pellets early in training and was later reduced to one pellet per visit.

#### Robot

Robots in cohort 1 and cohort 2 differed in their construction. Cohort 1 employed the LEGO Mindstorms SPIK3R set (set #31313, Billund, Denmark) assembled into a scorpion-like configuration ^1^. For cohort 2, we reconfigured the LEGO EV3 components into a cobra-inspired design and mounted them on an OSOYOO® omnidirectional chassis (Pinetree Electronics Ltd, Richmond, BC, Canada). To enhance the robot’s predator-like appearance, we affixed large “googly” eyes to its front. During an attack, the robot emitted a warning screech and lunged forward when triggered, with the attack event lasting approximately 2.8 seconds.

#### Experimental Timeline

Prior to surgery, cohort 1 rats were trained to run a minimum of 50 laps on the linear track for two consecutive days. Cohort 2 did not receive any pre-training before implantation.

Rats ran one 1-hour session per day. During LT, the robot bay was concealed and rats simply shuttled between feeders. LT sessions were repeated until rats reached a 50 lap threshold for cohort 1 and a 100 lap threshold for cohort 2. Cohort 1 then underwent a 2 day NOV phase in which the robot was present but non-attacking; cohort 2 omitted this phase. During the ATTK phase, the robot was triggered when the rat crossed an unmarked threshold. In both cohorts, the first 15 laps were safe. In cohort 1, attacks occurred with a 20% probability on each subsequent threshold crossing ^1^. In cohort 2, attacks were delivered according to a pseudo-random encounter schedule: the first attack occurred within the next five threshold crossings, and subsequent attacks occurred approximately once every 25 crossings. During the subsequent EXT phase, cohort 1 encountered a non-attacking robot, whereas cohort 2 had the robot bay closed and the robot fully removed. Finally, in the ReEX phase, both cohorts received an additional attack session under the same attack probability used during ATTK.

### Analyses

Behavioral events were identified from the linearized position signal using a common analysis pipeline applied to both cohorts. Although LT, NOV, and ATTK stages of cohort 1 were previously analyzed in Calvin et al.^1^, for the present manuscript cohort 1 was re-analyzed using the same event-detection and segmentation procedures as cohort 2 to ensure that the same pipelines were used for both cohorts. In this re-analysis, MTA-, MTC-, and retreat-related trajectories were segmented into distinct behavioral epochs (outbound, pause, inbound, and continued outbound (for MTCs only)).

#### Mid-track aborts (MTAs)

MTAs were defined as outbound excursions in which the animal exited the nest, advanced into the corridor, and then returned to the nest without reaching the far feeder. Candidate events were required to cross the doorway threshold and remain outside of the doorway for at least 2 seconds. To prevent attack-triggered retreat responses from being classified as MTAs, candidate MTAs were excluded if the excursion peak occurred within 2 s after a robot attack. For each detected MTA, the trajectory was segmented into three epochs: 1) an outbound segment extending from the doorway threshold to the end of sustained outbound movement, 2) a pause segment extending from the end of sustained outbound movement to the onset of sustained inbound movement (lasting at least 1 s), and 3) an inbound segment extending from the onset of sustained inbound movement to the doorway threshold crossing.

#### Mid-track continues (MTCs)

MTCs were defined similarly to MTAs, except that following the pause the animal resumed forward movement toward the far feeder rather than aborting and returning to the nest. Within each outbound journey, pauses were identified as gaps of at least 1 s between two sustained outbound movement bouts. As with MTAs, candidate events were excluded if the pause peak occurred within 2 s after an attack. MTCs were segmented into three epochs: 1) outbound (same as MTAs) and 2) pause (same as MTAs) and 3) a second outbound trajectory extending from pause offset to arrival at the far feeder.

#### Retreats

Retreats were defined as attack-triggered return trajectories. For each attack, the first MTA-like event occurring within 2 s after an attack was identified as the retreat. Retreats too were then segmented into three epochs: 1) outbound (defined identically to above), 2) post attack, extending from attack onset to retreat peak, and 3) inbound (again defined as above).

#### Control journeys

To provide baseline comparisons for each behavioral segment, control journeys were generated from the same session and corresponding movement direction. Control windows were selected from laps that did not contain attacks and were required not to overlap with any real event windows (including MTA, MTC, or retreat segments). For each event window, the corresponding control was chosen by matching the start and end positions of the event along the linearized track within a lap of the same direction.

### Neural Recording and preprocessing

#### Neural Recordings

Neural activity was recorded using chronically implanted silicon probes targeting the dorsal hippocampus, as described above. Signals were acquired using an Intan (cohort 1) or OpenEphys (cohort 2) recording system and digitized at 30 kHz. Spike and LFP signals were extracted from the raw recordings using standard filtering procedures. Spike data were obtained from high-pass filtered signals, whereas LFPs were obtained by low-pass filtering and downsampling the broadband signal to 2000 Hz for subsequent spectral analyses. Line noise was attenuated using a 60 Hz notch filter. Recordings were referenced using the acquisition system’s default referencing scheme, and channels with excessive noise or instability were excluded from further analysis.

During the task, cohort 2 additionally received closed-loop electrical stimulation of the hippocampal commissure during day 1 or 2 (counterbalanced) of ATTK as part of a separate experimental manipulation. Because this manipulation did not produce measurable effects on behavior or hippocampal signals relevant to the present analyses, data from stimulated and non-stimulated sessions were pooled.

#### Cell Identification

Previously sorted datasets from Calvin et al.^1^ were used for LT, NOV, and ATTK. For new data, spike sorting was performed using Kilosort (version 2) followed by manual curation in Phy. Previous unsorted cohort 1 data for EXT and ReEX and all data for cohort 2 were processed and curated using the same pipeline. Units were included if they were well-isolated based on waveform characteristics and cluster separation and exhibited stable firing throughout the recording session. Units with clear refractory periods and minimal inter-spike interval violations were retained. To ensure reliable decoding, sessions were excluded if fewer than 10 simultaneously recorded units were available.

#### Decoding

Spatial decoding was performed using a one-step Bayesian decoder based on ensemble spiking activity and occupancy-normalized spatial tuning curves derived from the animal’s linearized position. Spike trains from all simultaneously recorded units were converted into time-binned spike-count matrices. Spatial tuning curves were computed by dividing spike counts by occupancy within spatial bins of the linearized track. Decoded posterior probability over position P(x∣s) was estimated from the ensemble spike counts assuming independent Poisson firing across cells and a uniform prior over position.

For event-aligned analyses, decoded posterior probabilities were aligned to behavioral event anchors using peri-event time histograms. Unless otherwise specified, decoding was evaluated over a -2 to 2 s window surrounding the event using 100 ms time bins. For visualization, decoded posterior probabilities were first averaged across events within each session so that each session contributed one peri-event decoding estimate to the group summary. The animal’s linearized position was aligned to the same events and overlaid as the mean ± SEM trajectory.

#### Warping (Position and Time)

To compare behavioral epochs that differed in absolute duration or trajectory length, event-aligned decoding was normalized to a common progress axis before averaging across events. Warped decoding was performed after Bayesian decoding had first been computed in real time for each event window.

For pause epochs, decoding was *warped in time.* For each event, posterior probability over position and the animal’s linearized position were restricted to the event window and then interpolated onto a normalized time axis spanning 0 to 1, where 0 corresponded to event onset and 1 corresponded to event offset. This procedure preserved the relative temporal progression of the pause while allowing pauses of different durations to be directly compared. Events shorter than the minimum pause duration (1 s) were excluded.

For outbound and inbound movement epochs, decoding was *warped in position*. Posterior probability and linearized position were first sampled across each event window, and the event trajectory was then transformed to a normalized position-progress axis spanning 0 to 1, where 0 corresponded to the start of the trajectory and 1 corresponded to its end. To reduce the influence of small tracking fluctuations or brief reversals in movement, position progress was constrained to increase monotonically prior to interpolation

In both warped-time and warped-position analyses, events were represented using 30 normalized bins. Warped decoding was computed separately for the two behaviors being compared within each session, and session-level decoding-difference maps were then calculated as the mean decoded posterior during condition A minus condition B.

#### Region Definitions

For region-based analyses, the linearized track was divided into five predefined regions: Nest, Track Start, Attack, Robot, and Far Feeder. Region boundaries were derived from the linearized positions of the nest feeder, doorway, attack threshold, robot, and far feeder. Specifically, the Nest region extended from the nest feeder to the doorway, the Track Start region from the doorway to the midpoint between the doorway and attack threshold, the Attack region from that midpoint to the midpoint between the attack threshold and robot, the Robot region from the attack-robot midpoint to the midpoint between the robot and far feeder, and the Far Feeder region from the robot-far feeder midpoint to the far feeder.

#### Quantification of Decoding Differences

For each session, decoding-difference maps were averaged across all normalized progress bins and all spatial bins within each track region, yielding one mean decoding-difference value per region per session. Positive values indicate stronger representation during the first condition in the subtraction, whereas negative values indicate stronger representation during the second condition.

#### Power spectral density analyses

PSD analyses were performed on local field potentials decimated from the full 30 kHz recordings to 2 kHz. Spectral power was estimated using Fourier transform-based spectrograms computed from the full LFP trace using a 250 ms sliding window, evaluating frequencies from 1-150 Hz and 1-25 Hz for the inset figures.

Event-related spectra were obtained by aligning spectrograms to behavioral event anchors and extracting spectral power within a fixed time window relative to the alignment point. PSDs were aligned to the start of the relevant segment and computed over the 0-1 s interval following alignment.

For each event, spectral power within the analysis window was averaged across time bins at each frequency and log10-transformed. Spectra were first averaged across events within each session, yielding one mean spectrum per session for each event type. Group summaries were then computed across sessions and are reported as mean ± SEM across sessions.

For event-aligned spectra, spectrograms were aligned to event anchor times using peri-event time histograms. For each event, spectral power was extracted within a peri-event window and averaged across events within session. Session-level peri-event spectra were then averaged across sessions for visualization. Spectral power is shown as log10-transformed power across frequency and time relative to the event.

### Statistical analysis

#### Behavioral analysis

Behavioral analyses were conducted at the session level unless otherwise noted. For each rat and session, we quantified lap count, MTA count, MTC count, retreat count, and mean inbound and outbound speed. Mixed-effects models ^47^ were used to account for repeated measurements by including rat identity as a random factor.

Continuous outcomes (lap counts and normalized speeds) were analyzed using linear mixed-effects models, whereas count data (MTA and MTC events) were analyzed using Poisson generalized linear mixed-effects models. Models included phase as a fixed effect; analyses comparing cohorts additionally included cohort and the phase × cohort interaction. Planned comparisons tested differences between key phases (ATTK vs LT, EXT vs ATTK, and ReEX vs LT).

Abort probability and retreat proportion were analyzed as session-level proportions. Abort probability was defined as # of MTA/(# of MTA + # of MTC), and retreat proportion as # of retreats / # of attacks. Changes across attack sessions were tested using linear mixed-effects models with session number as a fixed effect, and differences between ATTK and ReEX were tested using phase as a fixed effect.

For inbound and outbound speed analyses, session-level mean speeds were normalized within rat to each animal’s baseline speed during the LT phase. Specifically, each value was divided by that rat’s mean LT speed for the corresponding direction, so that normalized values represent proportional change relative to the animal’s own linear-track baseline.

For Poisson generalized linear mixed-effects models, overdispersion was assessed using the Pearson chi-square statistic divided by residual degrees of freedom. Degrees of freedom for linear mixed-effects models were estimated using the Satterthwaite method.

#### Decoding Analyses

To quantify differences in hippocampal representations between behavioral events, decoding difference maps were computed as the mean decoded position during one event minus the other (e.g., Retreat Inbound - MTA Inbound; Fig. 3E). For each session, decoding differences were averaged within the five track regions described above, yielding one mean decoding-difference value per region per session.

Region-level effects were assessed using a nested analysis of variance (ANOVA) with region and experimenter as fixed effects and rat nested within experimenter as a random effect. Post-hoc comparisons tested whether decoding differences within each region differed from zero using Wilcoxon signed-rank tests across sessions. Bonferroni correction was applied to control for multiple comparisons across the five regions.

### Computational Resources

All analyses were performed in MATLAB (version R2024b; MathWorks, Natick, MA). Custom scripts were developed to perform behavioral segmentation, decoding, and spectral analyses. Data from cohort 1 were reanalyzed using this unified MATLAB-based pipeline to ensure consistency across cohorts.

